# Time-resolved molecular measurements reveal changes in astronauts during spaceflight

**DOI:** 10.1101/2023.03.17.530234

**Authors:** Minzhang Zheng, Jacqueline Charvat, Sara R. Zwart, Satish Mehta, Brian E. Crucian, Scott M. Smith, Jin He, Carlo Piermarocchi, George I. Mias

## Abstract

From the early days of spaceflight to current missions, astronauts continue to be exposed to multiple hazards that affect human health, including low gravity, high radiation, isolation during long-duration missions, a closed environment and distance from Earth. Their effects can lead to adverse physiological changes and necessitate countermeasure development and/or longitudinal monitoring. A time-resolved analysis of biological signals can detect and better characterize potential adverse events during spaceflight, ideally preventing them and maintaining astronauts’ wellness. Here we provide a time-resolved assessment of the impact of spaceflight on multiple astronauts (n=27) by studying multiple biochemical and immune measurements before, during, and after long-duration orbital spaceflight. We reveal space-associated changes of astronauts’ physiology on both the individual level and across astronauts, including associations with bone resorption and kidney function, as well as immune-system dysregulation.

## Introduction

As human space exploration continues to expand we anticipate an increasing number of long-duration and deep space missions. Beyond missions in low Earth orbit (LEO), plans are currently underway for a return to the moon, and utilizing such missions towards building a gateway for inter-planetary travel to Mars. Half a century of long-duration spaceflights on space stations has already demonstrated that such missions will pose several health risks to astronauts that have to be addressed. There are five categories of hazards faced by astronauts, that can directly contribute to health risks ^1,2^: (i) Transitioning from Earth’s gravity to effective weightlessness or lower-than-Earth gravity. (ii) Experiencing higher levels of radiation, with considerably less shielding in the absence of Earth’s protective magnetic field. (iii) Being in isolation during long-duration missions. (iv) Living in a closed environment with an evolving ecosystem for long periods of time. (v) Traveling and living at vast distances from Earth, where communications and facilities will be limited for direct health care assessment and intervention if necessary. The multiple physiological effects associated with these hazards include muscle and bone weakening, kidney stone risks and other problems due to fluid redistribution in the upper body, cancer risks due to radiation, immune dysregulation, cognitive and behavioral effects, and nutritional deficits ^2–4^. All these hazards and associated risks dynamically change during space missions and necessitate close monitoring of crew members’ health to ensure their safety and well-being.

Astronauts have experienced multiple clinical symptoms in long-duration LEO flights, with notable medical events reported in up to 46% of the crew members ^4^. The most frequently reported events (defined as involving symptoms that are recurring or have prolonged duration, or are unresponsive to treatment), included cold sores, rashes, allergies, and infectious diseases ^4^. Several studies aim to understand the molecular basis of these medical symptoms in long flights. Krieger et al. ^5^ investigated cytokines in plasma and saliva in 13 astronauts during 4-6 month International Space Station (ISS) flights, and identified persistent immune dysregulation. Nutritional deficits have also been directly linked to space-related clinical symptoms and systemic physiological changes ^6^. Smith et al. evaluated astronaut nutritional status during long stays on the ISS, identifying multiple marker changes associating bone resorption and oxidative damage across 11 astronauts ^7^. Given the mission-critical role of maintaining astronaut health, the National Aeronautics and Space Administration (NASA) has developed the integrated medical model (IMM) as a tool for identifying medical conditions and quantifying medical risks to astronauts ^4,8,9^. Furthermore, the monitoring of standard medical physiological measurements, immune and metabolite markers, is now being supplemented with an expanding array of thousands of molecular measurements (omics) in individual astronauts. This was exemplified by the astronaut Twins Study ^10^, which focused on molecular biology measurements to compare the effects of spaceflight on two twins, one on a 340-day mission, while the other was observed during this period on Earth. Multiple changes were observed giving a glimpse of how long-duration spaceflight affects an individual astronaut’s physiology.

The next steps must enable monitoring of data from multiple astronauts during long-duration space missions, and should include the implementation of a personalized approach that can establish baseline characteristics in each crew member, detect deviations therefrom, and detect potentially adverse medical events. Such personalized monitoring of health based on omics have been implemented in the NASA Twins Study ^10^, and on Earth, using blood ^11^, urine, saliva ^12^ and fecal samples. Testing methods have included microbiomes ^13^, personalized coaching ^14^ and digital devices ^15^. In this investigation, we implemented the first (to our knowledge) multi-astronaut individual-focused monitoring in long-duration spaceflight. We characterized personal biological signals in 27 astronauts (8 female, 19 male) and assessed multiple metabolites, immune cell constitution, and physiological measurements spanning long-duration orbital missions (mean flight duration *∼* 165 days), Figure 1. Using spectral-based methods ^12,16,17^ we identified temporal trends, and classified multi-signal responses corresponding to in-flight and return to Earth changes. Our analysis addressed typical spaceflight research limitations, including monitoring issues that include uneven time sampling and missing data, as well as different sampling rates for different astronauts. We identified a set of time-resolved changes in metabolites and immune cell behavior that was concordant across the majority of astronauts, and associated with bone resorption, renal stone formation, T cell and monocyte dysregulation and nutritional deficiencies. Finally, we constructed a network assessing inter-astronaut similarities, identifying groups of astronauts with similar time-resolved trends and reported clinical symptoms. Our findings identify significant metabolite and immunological modifications during spaceflight, and suggest that individual-focused monitoring must be implemented to identify the onset of these changes in order to also address them in a timely manner with personalized countermeasures.

**Fig. 1.**
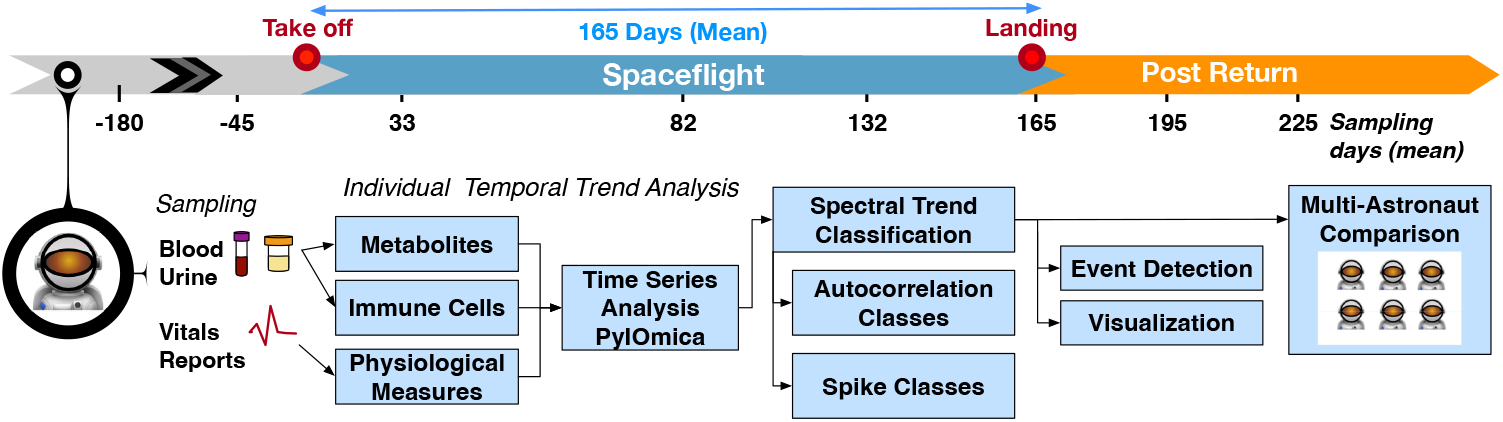
Study overview. The study followed 27 astronauts across multiple long-duration missions (adjusted mean 165 days) on ISS. The data included multiple time measurements from 180 days pre-flight to two months post-flight for different blood and urine-detected molecules, as well immune cell characterizations, physiological measurements, and medical reports. The time-resolved measurements were analyzed per astronaut to identify temporal trends and detect events of interest. The individual measurements were finally used to construct a multi-astronaut comparison based on similar temporal changes across measurements.

## Results

### Astronaut cohort

We investigated time-resolved measurements from NASA astronauts across multiple missions to ISS. These astronauts had participated in two studies: the Nutritional Status Assessment and Validation of Procedures for Monitoring Crewmember Immune Function. These studies included multiple metabolic and physiological measurements (264 annotations), including blood and urine metabolite analyses and immune cellular assessments which were taken over multiple time-points before, during, and after flight.

This was a retrospective analysis of existing data. As part of our broader study on astronaut integrative data analysis, we identified 47 astronauts meeting the criteria of having multiple time-resolved measurements, and of those, 38 astronauts agreed to participate in our study following informed consent for this additional analysis. They had previously provided informed consent to participate in the original studies mentioned above. Out of these, 27 were found to have long-duration mission data (165 days mean) and were selected for analysis. Data were obtained for the Life Surveillance for Astronaut Health (LSAH) and Life Sciences Data Archive (LSDA).

Over the course of their missions, the astronauts experienced several clinical symptoms that were provided to our team as IMM reports. We aggregated the information across astronauts and events in Table 1. Events that resolved within the first 30 days of flight are classified as space adaptation symptoms ^4^. The top space adaptation events included space motion sickness (14 astronauts), headache (10), and insomnia (9). The most frequent events later in the mission, categorized as non-adaptation events, included sleep disorder (15; reported 34 times), fatigue (14), and skin rash (9).

**Table 1.**
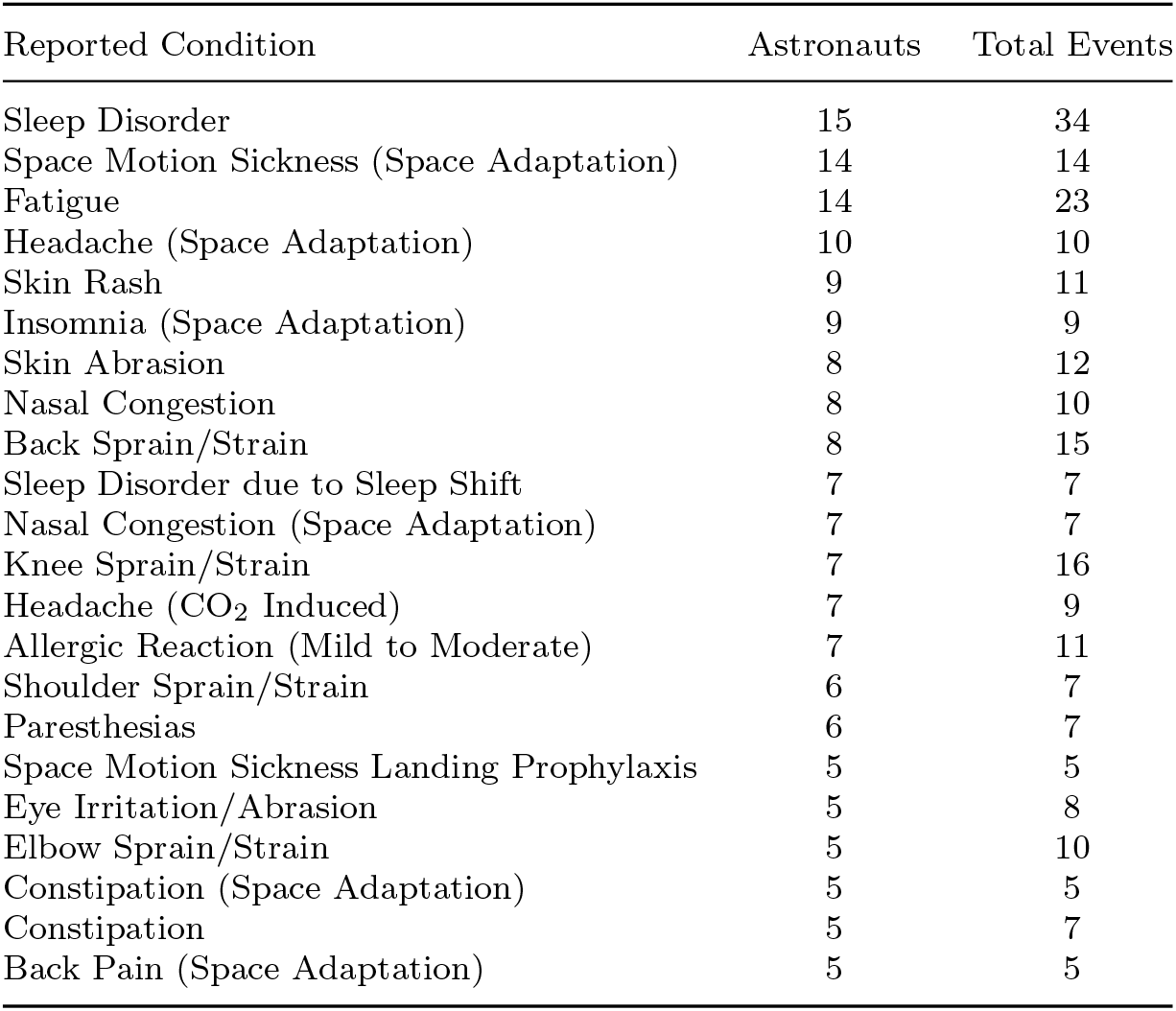
Integrated Medical Model Reports

As is common, not only in spaceflight but with any medical monitoring outside laboratory settings, different measurements were available for different astronauts, with highly heterogeneous and uneven time sampling, and missing data. We created a common time sampling grid across all astronauts, where measurement data for each astronaut were summarized by time averages into nine adjusted mean-based timepoints: 180 and 45 days prior to flight, early flight (33 days), mid-flight (82 days) and late flight (132 days), return to Earth (after 165 days of flight), and 30 and 60 days following return. The common time grid was used to: (i) ensure non-attributability of data back to individual astronauts, since the time of flight is sometimes unique to each astronaut, and to enable the individual measurements to be comparable across astronauts.

### Individual astronaut time-resolved assessment

We classified each astronaut’s time-resolved measurement into different classes using spectral methods ^12,16,18^ implemented in `PyIOmica` ^17^. The approach allowed the identification of sets of measurements that display time-resolved deviations from an astronaut’s baseline. This categorization provided 3 classes of time trends: (i) *Lags*, for measurements showing statistically significant auto-correlation at different lags. Here an autocorrelation of a signal at a certain lag refers to a correlation of a signal with a delayed (lagged) version of itself. (ii-iii) *Spike Maxima or Minima* for measurements that do not belong in the *Lags* class, but have punctuated intensities (spikes) that are either high (Maxima) or low (Minima) at particular timepoints. The measurements assigned to these classes are further clustered into groups (G) and subgroups (S) based on their overall autocorrelation and signal intensity similarities respectively (see Methods).

The individual astronaut signal analyses are available in the Online Data Files (ODFs). Examples of the analysis are shown in Figure 2 for two astronauts. The heatmaps include the intensities of the measurements that had statistically significant (FDR *<* 0.05) trends for the Lag 1 class in two astronauts. Clustering of the data into groups (G) reveals similarities due to the signal autocorrelations, with the corresponding autocorrelation heatmap depicted on the right of the figure. The heatmap on the left depicts the intensities of the measurements and the final subgroup structure. For each subgroup, a summary of the time-resolved median structure is summarized with a graphical representation by a visibility graph ^19^.

**Fig. 2.**
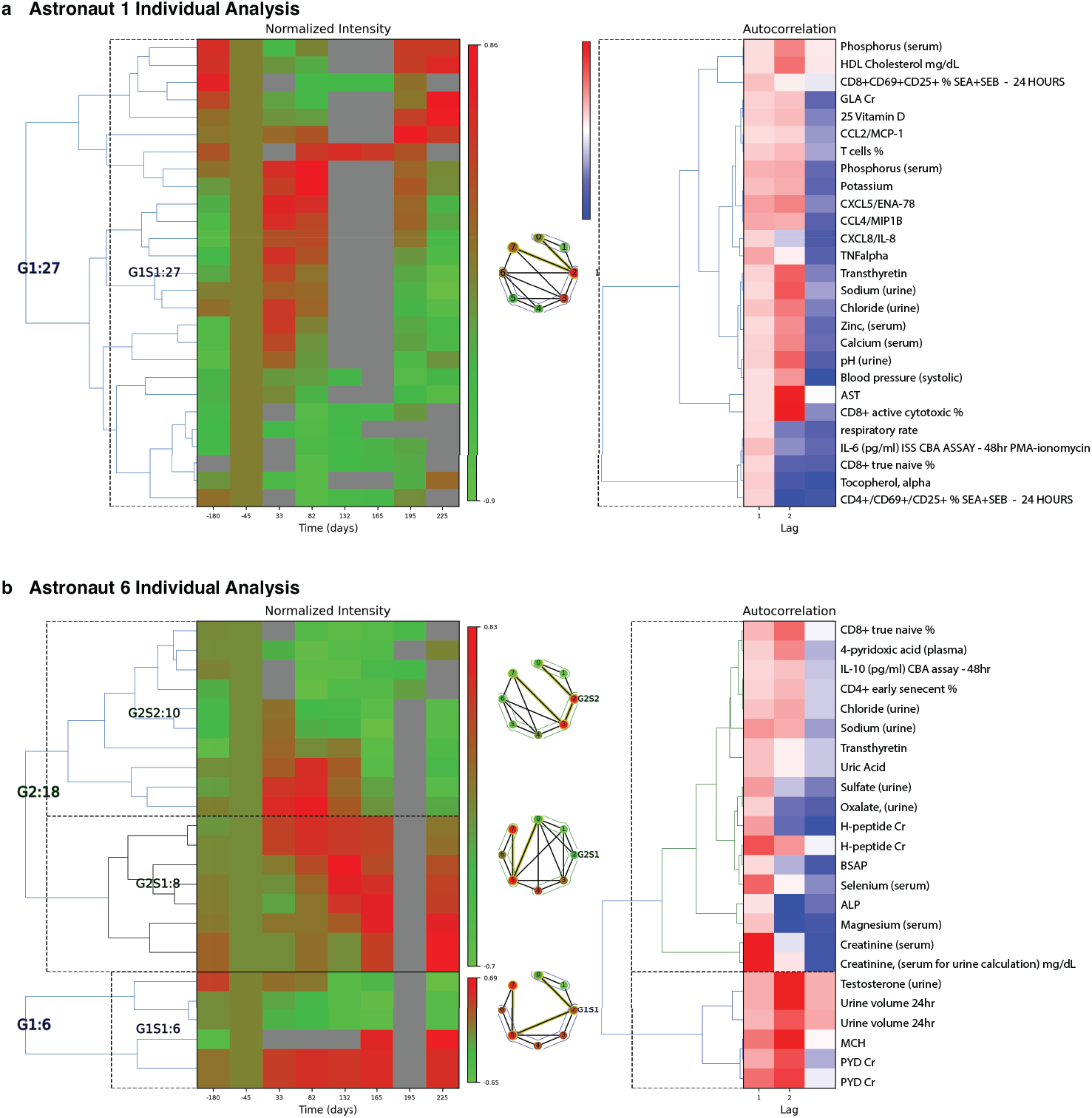
Individual Astronaut Examples. Classification results (Lag 1 patterns) are shown for astronaut 1 (a) and 6 (b). For each astronaut, we have: *Left heatmap*: Groups/sub-groups clustering of measurements, with adjusted mean time (with respect to takeoff). Times 165, 196 and 225 represent return to Earth, one and two months post-return respectively. *Middle graph*: Time-resolved visibility graph representation of median behavior within each group, with visibility-graph community detection identifying similar time behaviors ^19^. *Right heatmap*: Autocorrelation clusters and annotations for the measurements identified. *ALP: alkaline phosphatase, BSAP: bone-specific alkaline phosphatase, CBA: cytometric bead assay, Cr: creatinine normalized, G: group, GLA: gamma-carboxyglutamic acid, MCH: mean corpuscular hemoglobin, PMA: Phorbol 12-myristate 13-acetate, PYD: pyridinium cross-links, S: subgroup, SEA: staph enterotoxin A, SEB: stap enterotoxin B. Multiple evaluations/repeats included*.

For astronaut 1, Figure 2a, there are changes associated with Lag 1 class. The main trend consists of upregulation of a subset of immune markers early in flight (Days 33 and 82), and a return to pre-flight levels after return to Earth. The changes are associated with with plasma concentrations of the cytokine CXCL5, CCL4, CXCL8, and TNF alpha, indicative of immune dysregulation (potential inflammation). Additionally, in urine there is an increase in sodium, chloride and zinc in early flight, returning to post-flight levels upon return to Earth. *γ*-carboxyglutamic acid (GLA) is decreased during flight, returning to pre-flight conditioning levels two months post return. GLA reflects posttranslational modification of glutamic acid, and this modification is vitamin K dependent ^20^. On flights to the Russian space station Mir, vitamin K status in one crewmember was low during flight ^21^ and subsequently vitamin K supplementation has been offered as a countermeasure for bone loss in astronauts. On ISS, this was not observed in a larger cohort of astronauts on longer missions ^22^. In parallel, there is a decrease in immune cell levels, including CD8+ active and naïve relative percentages. This can be indicative of decreased immune response, as well as disruption of immune cell maturation cycles ^23^.

For astronaut 6 there are 3 sets of temporal patterns, Figure 2b, with changes associated with the Lag 1 class. In the first Group, MCH (mean corpuscular hemoglobin) and PYD (pyridinium cross-links) were found to increase during flight and stay high even following 2 months post return to earth. Increased MCH is associated with macrocytic anaemia. PYD is a bone resorption marker reflecting changes in bone metabolism and bone remodeling. With a symmetrically opposite trend, urine volume and testosterone decrease in flight and remain low for up to 2 months after return to Earth. The second set G2S1 indicates a gradual increase in intensities of H-peptide (another bone resorption marker), total and bone-specific alkaline phosphatase (BSAP), selenium, magnesium, and creatinine that persist after return to Earth. Probable indications include bone disease and kidney dysfunction, though certain values may be affected by dietary intake (e.g. magnesium). Finally, the third trend, G2S2 involves sets that show an increase in early flight but then consistent decreases in intensity that persist post flight. These include CD8+ T cell ‘true naïve’ and CD4+ T cell ‘early senescent’ relative percentages, that may indicate immune cell maturation dysregulation. IL-10 also remains reduced post flight, as compared to preflight baseline concentrations. Other compounds with similar trends include 4-pyridoxic acid, chloride and sodium in urine, transthyretin, uric acid, and sulfate and oxalate levels (in urine).

The results for all astronauts showed similar patterns, indicative of individual-specific changes particularly associated with early spaceflight or landing stress, with multiple changes persisting up to two months following return to Earth. Further details and representations of the trends identified for each astronaut are available in deidentified plots and tables in the ODFs.

### Spaceflight effects across multiple astronauts

To identify similarities across astronauts, we investigated how often the individual-astronaut statistically significant signals were identified across multiple astronauts. The top measurements by incidence (*≥* 16), are shown in Table 2, including a trend summary and potential physiological aspects.

**Table 2.**
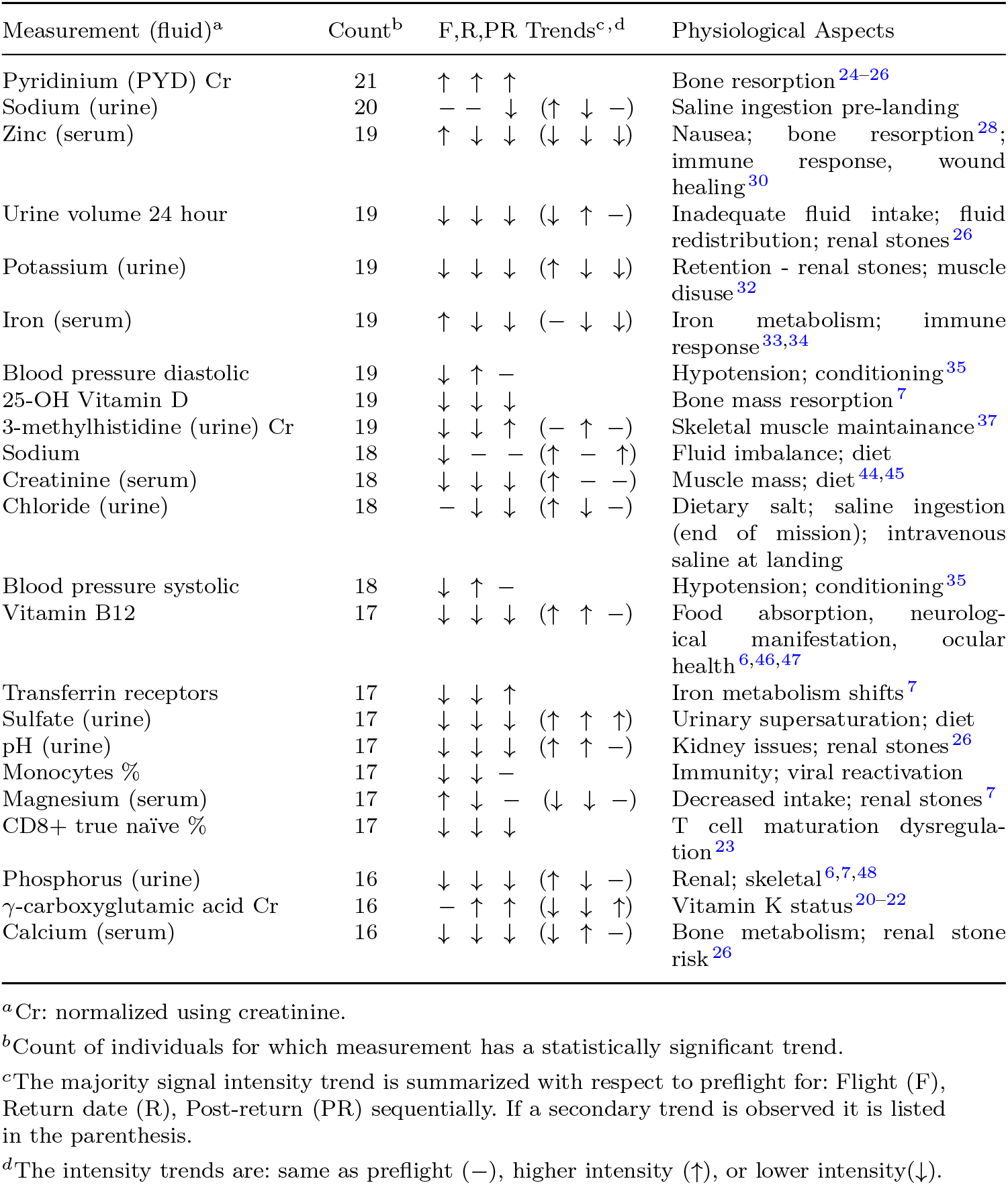
Aggregated Measurements Across Astronauts.

| Measurement (fluid)<sup>a</sup> | Count<sup>b</sup> | F,R,PR Trends<sup>c,d</sup> | Physiological Aspects |
| --- | --- | --- | --- |
| Pyridinium (PYD) Cr | 21 | ↑ ↑ ↑ | Bone resorption<sup>24–26</sup> |
| Sodium (urine) | 20 | – – ↓ (↑ ↓ –) | Saline ingestion pre-landing |
| Zinc (serum) | 19 | ↑ ↓ ↓ (↓ ↓ ↓) | Nausea; bone resorption<sup>28</sup>; immune response, wound healing<sup>30</sup> |
| Urine volume 24 hour | 19 | ↓ ↓ ↓ (↓ ↑ –) | Inadequate fluid intake; fluid redistribution; renal stones<sup>26</sup> |
| Potassium (urine) | 19 | ↓ ↓ ↓ (↑ ↓ ↓) | Retention - renal stones; muscle disuse<sup>32</sup> |
| Iron (serum) | 19 | ↑ ↓ ↓ (– ↓ ↓) | Iron metabolism; immune response<sup>33,34</sup> |
| Blood pressure diastolic | 19 | ↓ ↑ – | Hypotension; conditioning<sup>35</sup> |
| 25-OH Vitamin D | 19 | ↓ ↓ ↓ | Bone mass resorption<sup>7</sup> |
| 3-methylhistidine (urine) Cr | 19 | ↓ ↓ ↑ (– ↑ –) | Skeletal muscle maintainance<sup>37</sup> |
| Sodium | 18 | ↓ – – (↑ – ↑) | Fluid imbalance; diet |
| Creatinine (serum) | 18 | ↓ ↓ ↓ (↑ – –) | Muscle mass; diet<sup>44,45</sup> |
| Chloride (urine) | 18 | – ↓ ↓ (↑ ↓ –) | Dietary salt; saline ingestion (end of mission); intravenous saline at landing |
| Blood pressure systolic | 18 | ↓ ↑ – | Hypotension; conditioning<sup>35</sup> |
| Vitamin B12 | 17 | ↓ ↓ ↓ (↑ ↑ –) | Food absorption, neurological manifestation, ocular health<sup>6,46,47</sup> |
| Transferrin receptors | 17 | ↓ ↓ ↑ | Iron metabolism shifts<sup>7</sup> |
| Sulfate (urine) | 17 | ↓ ↓ ↓ (↑ ↑ ↑) | Urinary supersaturation; diet |
| pH (urine) | 17 | ↓ ↓ ↓ (↑ ↑ –) | Kidney issues; renal stones<sup>26</sup> |
| Monocytes % | 17 | ↓ ↓ – | Immunity; viral reactivation |
| Magnesium (serum) | 17 | ↑ ↓ – (↓ ↓ –) | Decreased intake; renal stones<sup>7</sup> |
| CD8+ true naïve % | 17 | ↓ ↓ ↓ | T cell maturation dysregulation<sup>23</sup> |
| Phosphorus (urine) | 16 | ↓ ↓ ↓ (↑ ↓ –) | Renal; skeletal<sup>6,7,48</sup> |
| γ-carboxyglutamic acid Cr | 16 | – ↑ ↑ (↓ ↓ ↑) | Vitamin K status<sup>20–22</sup> |
| Calcium (serum) | 16 | ↓ ↓ ↓ (↓ ↑ –) | Bone metabolism; renal stone risk<sup>26</sup> |

<sup>a</sup>Cr: normalized using creatinine.<sup>b</sup>Count of individuals for which measurement has a statistically significant trend.<sup>c</sup>The majority signal intensity trend is summarized with respect to preflight for: Flight (F), Return date (R), Post-return (PR) sequentially. If a secondary trend is observed it is listed in the parenthesis.<sup>d</sup>The intensity trends are: same as preflight (–), higher intensity (↑), or lower intensity(↓).

Multiple signals showed similar trends across individuals. The top examples are shown in Figure 3a-i across all measurements, as well as for immune cells, Figure 3j-l. The results from PYD are prominent and most prevalent (observed in 21 astronauts), Figure 3a, showing an increase in intensity beginning early in spaceflight, and also persisting post flight (at least for 30 days). PYD is a marker of bone resorption and is indicative of the ongoing concern of bone loss among astronauts on long-duration spaceflights ^7,24–27^. We also observe fluctuations and a decrease in urinary sodium, Figure 3a, that indicates retention particularly upon return to Earth, and this may be related to the effects of microgravity on body fluid balance, as also seen in urine volume Figure 3d.

**Fig. 3.**
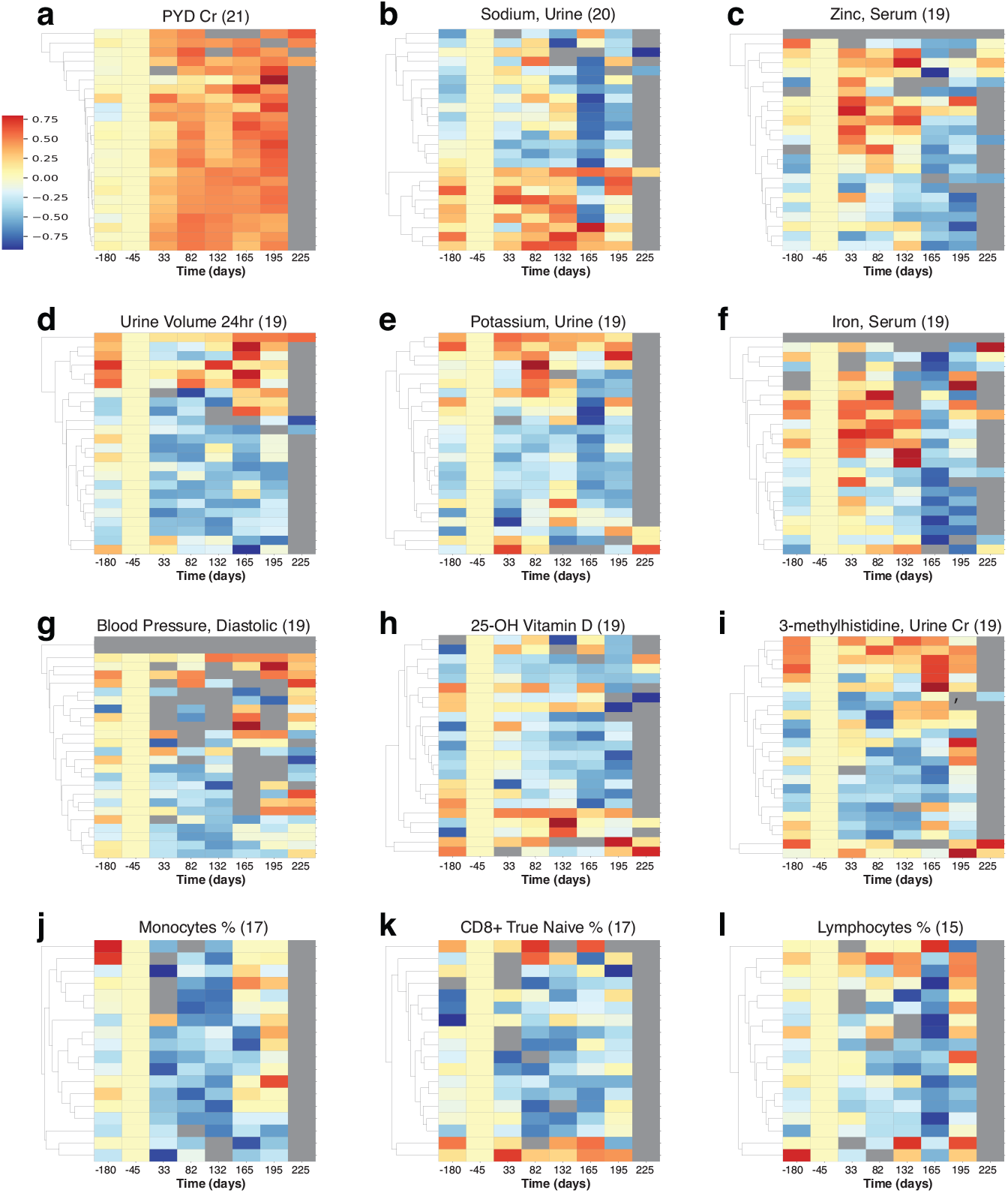
Multiple measurements show similar trends across astronauts. The heatmaps represent the normalized intensities of each measurement across all astronauts (rows) over time (columns). These measurements were identified to show statistically significant trends (FDR *<* 0.05) in multiple crew members, with the respective astronaut tallies shown in parentheses. a-i are the top measurements by frequency overall, and j-l are the top immune cell measurements. *PYD: pyridinium cross links, Cr: normalized with creatinine*.

Zinc in serum, Figure 3c, is elevated during flight, and returns to low levels after return to Earth ^28^. While zinc serum is not an indicator of nutritional zinc levels ^29^, the fluctuations may have health implications, as long term elevated zinc can induce nausea and headaches and gastrointestinal problems, potentially leading to decreased immune function and copper deficiency, while zinc deficiency could be an issue after return to Earth ^30^.

Potassium levels were found to be generally lower (but elevated for a small subset of astronauts during flight), with a sharp decrease across astronauts upon return to earth. This was also reported for short Apollo missions ^31^, and the connections between muscle loss, kidney dysfunction and cardiovascular conditions need to be investigated further ^32^.

Iron in serum, Figure 3f, appears affected by landing events as there is a marked decrease post return to Earth indicative of potential disruption of iron metabolism, which could lead to anemia and affect the immune response ^33^. There is also an increase in relative iron levels during flight for some astronauts, that may be related to neocytolysis ^34^. Diastolic blood pressure, Figure 3g, is reduced during space flight across astronauts; this transient in-flight effect has also been reported in a smaller cohort (n=12) ^35^ and could have implications for hypotension during flight (though returning to normal levels following Earth return). 25-OH Vitamin D measurements are decreased across 19 astronauts, consistent with concerns relating to bone health ^7^. 3-methylhistidine (3-MH), normalized with creatinine ^36^, was generally decreased during spaceflight, showing some increase in levels on landing for a subset of astronauts. An increase of 3-MH would be consistent with skeletal muscle degradation ^37^, and the overall 3-MH changes in flight, particularly punctuated on landing require further investigation.

In terms of immune cells, we noticed a general decrease in Figure 3j-l, in monocyte, CD8+ naïve T cell, and ‘bulk’ lymphocyte relative percentages.

This decrease highlights the potential for immune related disorders ^23,38,39^. This finding could also be linked the (largely) asymptomatic reactivation of latent herpesviruses, as has previously been observed in ISS astronauts during spaceflight^39^. It should be noted, that recently atopic dermatitis in some astronaut cases has been positively correlated with latent herpesvirus reactivation, in particular HSV1^40^. Recently, Mourad et al., 2022, postulated that reactivation of VZV occurred in mothers, who experienced an asymptomatic reactivation of herpes zoster virus, similar to that described in astronauts ^41–43^. Additionally, we investigated whether the common temporal trends across astronauts could provide a grouping of astronauts based on similar physiological responses, by studying the structure of a multi-astronaut network ^18^. In the constructed network, Figure 4, nodes depict astronauts (n=24) for which time series data were matched, while weighted edges represent how many measurements showed similarity (Euclidean distance) in their signal periodograms across pairs of astronauts. The five shown communities (labeled Community 0 though Community 4) were detected using a k-means approach, with k=5 (see Methods). While the number of astronauts included (n=24) restricts findings, we still observed community separations (except Community 4, which is a singleton), with distinct characteristics. The largest group, Community 0 included 7 male and 1 female astronaut. Weighted edges indicate that H Peptide drove the similarity within Community 0. During flight, the astronauts of Community 0 had reported space motion sickness predominantly (5 times) as well as sleep disorder, insomnia and headaches (4 times each). The second largest group (Community 1), is evenly balanced based on sex (3 male, 3 female). The clustering is dominated by MCV (mean corpuscular volume) behavior, and the members predominantly reported sleep disorder (5 times), as well as space motion sickness and skin rash (3 times each). Similarly, in Community 2 (3 male, 1 female) the astronauts reported insomnia and headaches (3 times each), while for Community 3 (3M, 2F) sleep disorder (4 times), and nasal congestion, shoulder sprain/strain and headache (3 times).

**Fig. 4.**
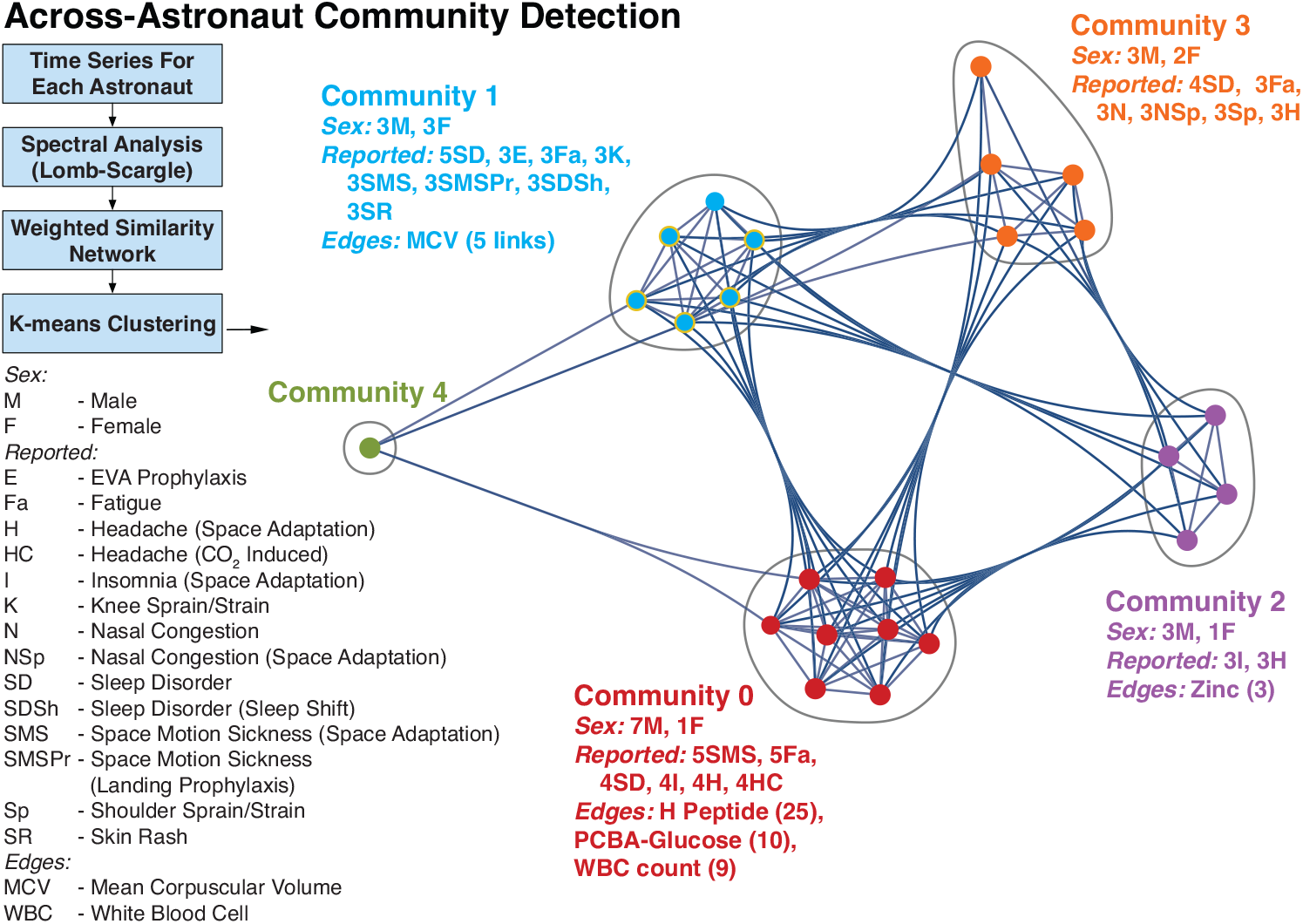
Astronaut Temporal Communities. Using the time-resolved measurements a network was constructed to identify communities of astronauts with similar temporal behavior. The nodes in the network represent astronauts, while the weighted edges correspond to different measurements, weighted with the number of astronaut pairs for which the time behavior is similar. Four Communities were identified (0-3) and a singleton, denoted in different colors. The male/female count, top reported IMM clinical symptoms, and edges (with corresponding counts/weights) associated with each community are included.

## Discussion

We have implemented an individual-focused approach to investigate personalized astronaut monitoring and detecting deviations from a wellness baseline. The astronauts experienced physiological changes that are associated with the identified molecular markers. The time-resolved analysis revealed the majority of changes to be associated with space stay, and also with the impact and stress of landing prior to readjustment to return to Earth. The impacted measurements are related to microgravity adaptation adverse effects such as bone resorption and kidney function, and to immune-system dysregulation. These changes suggest further countermeasures are necessary to prevent adverse outcomes (e.g. osteoporosis, renal stones, viral reactivation), particularly on long-duration flights. These can include enhanced food provisions, or if required, nutritional supplementation, as well as provision of relevant immune-disorder medications on board.

Our successful detection of individual-astronaut changes offers an extensible approach that can be used to integrate multiple time-resolved measurements (omics, physiological measurements, mobile device temporal data, and any temporal signal) on individual astronauts towards detecting and preventing potentially harmful medical events. By cataloguing such event onsets and associated measurement responses, the non-wellness transitions in astronauts can be used for providing timely diagnosis and countermeasures during space missions. This is vital during long-duration deep space missions, where distance from Earth necessitates self-reliance by astronaut crews to maintain their own wellness, as we expand space exploration to Mars and beyond.

## Acknowledgements

This project was supported by the Translational Research Institute for Space Health (TRISH) through NASA Cooperative Agreement NNX16AO69A (project T0412, PI: GI Mias). CP acknowledges support by NIH R01GM122085. The Nutritional Status Assessment SMO (PI: SM Smith) and the Validation of Procedures for Monitoring Crewmember Immune Function (PI: BE Crucian) were supported by the NASA Human Research Program Human Health Countermeasures Element.

## Author Contributions

Conceptualization, MZ, CP and GIM; Methodology, MZ, CP and GIM; Software, MZ and GIM; Investigation, MZ, JC, JH, SRZ, SM, BEC, SMS, CP and GIM; Visualization, MZ, GIM; Resources, MZ and GIM; Writing – Original Draft, GIM; Writing – Review & Editing, MZ, JC, JH, SRZ, SM, BEC, SMS, CP and GIM; Funding Acquisition, CP and GIM; Supervision, GIM.

## Methods

### Ethical approval

All protocols described herein were reviewed and approved by the NASA Institutional Review Board (NASA eIRB study ID: STUDY00000201) and by Michigan State University (MSU) Biomedical and Health Institutional Review Board (MSU study ID: STUDY00003581) in accordance with the requirements of the Code of Federal Regulations on the Protection of Human Subjects. Written informed consent was obtained from all participating subjects (astronauts). All methods described in this investigation were carried out in accordance with the relevant guidelines and regulations. Additionally, approvals for data use and manuscript/data submission for the study were obtained from the Lifetime Surveillance of Astronaut Health (LSAH) advisory board.

### Subjects

Forty-seven potential subjects were identified as U.S. astronauts that flew across 32 ISS and space shuttle expeditions and participated in two studies on Nutritional Status Assessment (SMO 016E), and Validation of Procedures for Monitoring Crewmember Immune Function (SMO 015). Thirty-eight astronauts agreed to participate following informed consent, and the analysis presented manuscript focuses on 27 subjects (20 male and 7 female) for whom long-duration measurements on the ISS were available (mean-based duration 165 days). Data for these subjects were obtained from LSAH and the Life Sciences Data Archive (LSDA).

### Data preprocessing

To create non-attributable (deidentified) data, for each astronaut the different measurements were summarized into a maximum of 8 time points (using averages of measurements within each time bin). These measurement time bins were labeled as -180 and -45 days for pre-flight; early flight (33 days), midflight (82 days) and late flight (132 days) measurements; return (165 days), return+30 days (195 days) and return + 60 days (225 days). The binned data were then used for two comparisons: (I) individual astronaut analysis and (II) and across-astronaut comparison.

### Individual astronaut analysis

#### Time series categorization

The data from each individual astronaut were analyzed using `PyIOmica`’s `calculateTimeSeriesCategorization` function ^17^. Here for each astronaut *a*, and each type of measurement *m*, the time series *X*_*ma*_ was analyzed, at time-points *t*_*i*_ *∈* {*−*180, *−*45, 33, 82, 132, 165, 195, 225} days. For each *X*_*ma*_ the difference in intensity of each timepoint *i* was computed relative to the pre-flight timepoint (−45 days) intensity, 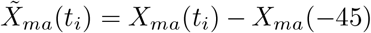. Finally, 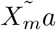 was normalized using Euclidean norm to *Q*_*ma*_, where 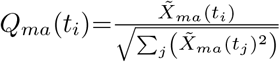.

The `calculateTimeSeriesCategorization` algorithm uses spectral methods to classify time series ^12,17^: Internally, the Lomb-Scargle periodogram *P*_*ma*_(*f*) of each time series is computed at a series of frequencies *f*. The inverse Fourier transform of *P*_*ma*_(*f*) yields the autocorrelations, {*ρ*_*mal*_ }, at lags *l∈ {*0, &, *N/*2}. A bootstrap time series set *B*_*ja*_(*t*), with *j∈*{ 1, &, 10^5^ }is also generated (by sampling with replacement across the time series of each astronaut). By computing the autocorrelations of each *B*_*ja*_(*t*), null distributions at each lag *l*, and the corresponding 0.95 quantiles {*ρ*_*ql*_ }are obtained. These quantiles are used to assign classes [*Lag M* ], where *M* is the smallest lag for which the autocorrelation at lag *l* of a signal *X*_*ma*_ is greater than the corresponding bootstrap distribution quantile:

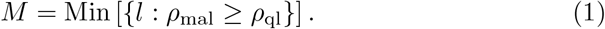

The algorithm also checks whether time series *X*_*ma*_ that do not fulfill the criteria in Equation (1) have any pronounced intensity maxima or minima. Using again a bootstrap set of time series, the 0.95 quantiles max_*qN*_ and min_*qN*_ are computed, from the distributions of the maxima and minima of the bootsrap time series respectively. If max(*X*_*ma*_) *>* max_*qN*_, *X*_*ma*_ is assigned to class [*SpikeMax*]. If min(*X*_*ma*_) *<* min_*qN*_, *X*_*ma*_ is assigned to class [*SpikeMin*]. Time series assigned to any of the classes [*LagM* ], [*SpikeMax*], and [*SpikeMin*] are considered to display statistically significant trends.

#### Cluster Categorization

For each time series class described above, `PyIOmica`’s `clusterTimeSeriesCategorization` was used to further resolve the time trends, by implementing two levels of agglomerative hierarchical clustering: (i) First-level group (G) clustering uses autocorrelations *ρ*_*mal*_ as a vector for computing similarities (correlation based) (ii) Second-level Subgroup (S) clustering uses the normalized time series *Q*_*ma*_ for computing similarities. The number of groups and subgroups within each class are determined by the silhouette method ^49^. For each astronaut and classification then results were summarized in heatmaps using `visualizeTimeSeriesCategorization` in PyIOmica. The [Lag 1] results from two astronauts are shown in Example outputs are shown in Figure 2. Full plots and corresponding measurements within each class are provided in the ODFs.

### Cross-comparison of astronaut time series

#### Astronaut graph-based communities

Graph-based community analysis of the astronaut time series was performed ^18^ with `networkx` ^50^ and `scikit-network`^51^. First, for each astronaut their time series periodograms *P*_*ma*_ were extracted using `PyIOmica`’s `LombScargle` method. A distance matrix with components [*D*_*m*_]_*xy*_ was computed, for each measurement *m* and astronauts *x, y*, where the entries correspond to the Euclidean distance between *P*_*mx*_ and *P*_*my*_. Again, a bootstrap set (50,000 signals) was constructed, and a distribution of distances between bootstrap time series was computed, with its 0.99th quantile *d*_*q*_. Then the restricted distance matrix *R*_*m*_ entries were constructed, so that distances within a radius *d*_*q*_ were set to 1, and otherwise set to zero:

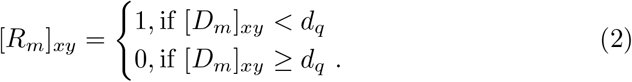

Using the restricted matrices for each measurement, a weighted adjacency matrix *A* = Σ _*m*_ *R*_*m*_ was obtained. The adjacency matrix represents a graph with astronauts as vertices and edges that connect astronauts. Hence, in the graph, an edge *E*_*xy*_ joins two astronauts *x, y* if there is at least one time series measurement *m* for which the pairwise distance between the periodograms *P*_*mx*_ and *P*_*my*_ was smaller than *d*_*q*_ (i.e. there is similar temporal behavior). Finally, the edge *E*_*xy*_ is given a weight that corresponds to the number of measurements for which there is temporal similarity (i.e. the number of measurements *m* with non-zero entries in the restricted distance matrix corresponding entry [*R*_*m*_]_*xy*_, Equation (2)).

#### Graph community detection

The community structure of the astronaut graph constructed above was determined: First, a graph embedding with generalized singular value decomposition (GSVD, dimension 2) of the adjacency matrix was carried out in `scikit-network`. The `sklearn.metrics.silhouette score` of ^52^ was used to determine the number of components based on silhouette scores ^49^. Finally, a consensus network was constructed where the procedure was repeated 1000 times, and nodes were allocated to their highest frequency community membership (highest probability assignment).

#### Code and Data Availability

All data files and code used in this investigation, referred to as Online Data Files (ODFs) in the manuscript, have been deposited to and are available from the NASA Life Sciences Data Archive (LSDA). LSDA is the repository for all human and animal research data, including that associated with this study. LSDA has a public-facing portal where data requests can be initiated (https://nlsp.nasa.gov/explore/lsdahome/datarequest). The LSDA team provides the appropriate processes, tools, and secure infrastructure for archival of experimental data and dissemination while complying with applicable rules, regulations, policies, and procedures governing the management and archival of sensitive data and information. The LSDA team enables data and information dissemination to the public or to authorized personnel either by providing public access to information or via an approved request process for information and data from the LSDA in accordance with NASA Human Research Program and Johnson Space Center (JSC) Institutional Review Board direction.

